# The role of inflammation in a humanized mouse model of transthyretin cardiac amyloidosis

**DOI:** 10.1101/2024.12.24.629308

**Authors:** Xiaokang Wu, Nixuan Cai, John Isaiah Jimenez, Hiroki Kitakata, Gracia Fahed, Alessandro Evangelisti, Alokkumar Jha, Joseph Woo, Ronglih Liao, Kevin M Alexander

**Affiliations:** Stanford Cardiovascular Institute, Stanford University, Stanford, CA, USA; Department of Medicine, Stanford University, Stanford, CA, USA; Neuroscience Graduate Program, Weill Cornell Medicine, New York, NY, USA; Memorial Sloan Kettering Cancer Center, New York, NY, USA; Cornell University, Weill Cornell Medical College, New York, NY, USA; Stanford University, Department of Cardiothoracic Surgery, Stanford, CA, USA; Stanford University, Stanford Amyloid Center, Stanford, CA, USA

**Author notes:** **Correspondence to Dr. Kevin M Alexander**,.

**Keywords:** Cardiomyopathy, Amyloidosis, Cardiac Amyloidosis, Transthyretin Cardiac Amyloidosis, TTR, ATTR, Animal Model, Inflammation

## Abstract

**Background:** Systemic amyloidosis represents a group of protein-misfolding diseases that confer significant morbidity and mortality for millions of patients worldwide. Transthyretin cardiac amyloidosis (ATTR) is a particularly devastating amyloid disease that affects middle-aged and elderly individuals and leads to cardiomyopathy (ATTR-CM), which has a median survival of 2.5 to 3.5 years [1, 2]. ATTR-CM can be hereditary, leading to a more aggressive disease course in younger patients. The most prevalent *TTR* variant in the United States is *V122I*, which is found in 3-4 % of African Americans [3]. Despite the significant healthcare burden, ATTR-CM remains underdiagnosed due to a lack of disease awareness and limited diagnostic techniques [4]. Informative *in vivo* models have proven elusive during the past decade [5]. Moreover, there is no available treatment to reverse cardiac dysfunction due to amyloid fibril deposition [1, 6, 7]. Therefore, a better understanding of the molecular mechanisms of ATTR-CM is imperative to developing novel, effective therapies.

**Method and Results:** To explore the pathogenesis of ATTR, we created a murine TTR knockout (TTR-KO) model expressing the human V122I *TTR* variant. To study the gender differences, both male and female TTR-KO mice were utilized in this study. Significant elevations of human TTR were observed in both male and female ATTR murine plasma post-injection 3 months (human TTR level (ng/ml) Male ATTR: 109.9 ± 5.568; Male control: 28.17 ± 7.010; p=0.0008, N=3 mice/group; Female ATTR: 127.5 ± 32.43; Female control: 20.08 ± 8.351; p=0.0327, N=3 mice/group) with preserved cardiac function (FS% Male ATTR: 26.07 ± 3.667; Male control: 22.69 ± 1.585; p=0.3712, N=6-8 mice/group; Female ATTR: 26.62 ± 1.980; Female control: 31.25 ± 4.482;p=0.3397, N=5-6 mice/group). Notably, the mouse model exhibited cardiac amyloid deposits confirmed by amyloidotic-specific Congo Red staining and Thioflavin T Staining. Transmission electron microscopy revealed both immature and mature amyloid fibrils in the extracellular matrix. RNA-sequencing of the ATTR mouse heart identified distinct transcriptomic patterns and conserved inflammation pathways similar to those seen in a cohort of human ATTR heart samples, including leukocyte transendothelial migration, T-cell receptor signaling, and apoptosis, along with upregulation of inflammatory markers CXCL-1/2/3 and CCL20, were observed in ATTR murine hearts. At the posttranslational level, we confirmed an increased level of CCL5 (MFI ATTR: 801 ± 105; Control: 426± 64; p=0.0061, N=3 mice/group) in murine plasma post-injection 3 months by a luminance-based immunoassay. The CXCL- and CCL-chemokines family are critical for directing leukocytes to inflammation sites.

**Conclusion:** In this study, we developed a humanized V122I ATTR mouse model with elevated circulating human TTR level and Congophilic amyloid deposits in the murine heart and kidneys. Our transcriptomic study suggested that inflammation may contribute to the ATTR-CM pathogenesis. Further studies are needed to decipher the precise interactions between inflammation and ATTR-CM.

**Highlights/What’s new/Clinical relevance:**

- We developed a humanized mouse model to replicate the multisystem complexity and clinical diversity associated with V122I ATTR-CM.
- Our study unveiled the pathogenic molecular mechanisms of amyloid deposition in ATTR-CM via a novel mouse model.
- We identified signature inflammatory pathways that uncover potential therapeutic targets for ATTR-CM.
- Our ATTR mouse model allows for preclinical pharmacogenomic assessments of novel therapeutics, which will undoubtedly improve outcomes for ATTR-CM patients.

## Introduction

Amyloidosis is a multisystemic disease in which proteins with unstable structures misfold and aggregate into amyloid fibrils and deposit in the heart and other organs [8]. The most common amyloid fibril proteins that can infiltrate the heart and lead to cardiac amyloidosis are immunoglobulin light chain amyloid fibril protein (AL) and transthyretin (TTR) amyloid fibril protein [9–12]. The TTR protein is mainly produced by the liver and is responsible for the transport of thyroid hormone and vitamin A. When TTR protein dissociates from native tetramer into monomers and misfolds, consequences aggregate to form oligomers, protofilaments, and mature amyloid fibrils which will deposit extracellularly in interstitial space of many organs causing transthyretin amyloidosis (ATTR), particularly the nervous and cardiac system [1, 10, 13]. Transthyretin amyloid cardiomyopathy (ATTR-CM) is a life-threatening, progressive, infiltrative disease that is often overlooked as a cause of heart failure, which confers significant morbidity and mortality leading to heart failure, arrhythmia, and sudden death [2, 6, 14].

Hereditary ATTR-CM patients often are heterozygous for a point mutation leading to an amino acid substitution in TTR. These mutations can also destabilize TTR tetramers and promote dissociation. To date, over 120 TTR variants have been identified as a cause of hereditary ATTR. In the United States, the most predominant TTR mutation is V122I, which is the TTR mutation that substitutes valine for isoleucine at position 122 (V122I) [15–17]. There are an estimated 3-4% of African Americans with 1.5 million people in the country carrying the V122I allele. Importantly, V122I TTR is associated with a 47% increased risk of heart failure and in a recent study was the 4th most common cause of heart failure among African descendants, which significantly increases the risk of developing hereditary ATTR-CM [3, 6, 18, 19]. V122I ATTR-CM was found to be linked to cardiac TTR amyloid accumulation in all carriers after the age of 65. Another major type of ATTR-CM is the wild-type (wtATTR-CM) which is thought to be due to age-related changes in the stability of TTR proteins, which predominantly affect older, Caucasian men [7, 20]. Once diagnosed, untreated patients with ATTR-CM have a median survival of 2.5 to 3.5 years [1, 2, 20]. However, a lack of disease awareness and limited well-established amyloid centers in the states, ATTR-CM diagnosis is frequently delayed or overlooked [6, 9, 21, 22]. Though it used to be considered rare overall, it is much more prevalent than previously thought with an estimated 13% of patients hospitalized with heart failure with preserved ejection fraction (HFpEF) and 16% of patients undergoing transcatheter aortic valve replacement (TAVR) having wtATTR-CM [23, 24].

Little is known about the cause of ATTR-CM and the lack of preclinical models hampers the discovery of therapeutics [5, 25–27]. There is currently no effective treatment to eliminate end-organ damage due to cardiac amyloid deposition, making its prevention and early diagnosis critically important [21, 28–30]. Many patients with end-stage ATTR will require a heart transplant, yet there is a profound shortage of donors [23, 31–33]. Thus, there is an unmet need to determine the cause and underlying molecular mechanisms that regulate ATTR pathogenesis and develop effective therapeutic targets for this devasting disease.

It is widely known that after cardiac injury, immune responders, such as neutrophils and macrophages, are the mediators of cardiac inflammation and subsequent repair [34–36]. Previous studies have shown that ATTR deposits in patients activate inflammatory and oxidative stress pathways [37]. A proteomic analysis from the Mayo Clinic was one of the first to identify a signature ATTR proteomic profile which revealed an association between high levels of PIK3C3 (a protein with a central role in autophagy) and worse survival in ATTR-CM patients [38]. Both macrophages and fibroblasts have been described to co-localize with TTR in the skin and gastrointestinal tract of TTR V30M mice [39]. The latest study showed a clear tendency of higher macrophage convergence to the TTR deposition sites in an HSF ± hTTR V30M mouse model of ATTR [40]. These data suggest a potential link between inflammation and the progression of ATTR-CM. However, the precise cellular and molecular signature inflammatory pathways involved in ATTR amyloidosis remain poorly understood.

This study introduced a novel humanized V122I ATTR mouse model onto a murine TTR-null background to address the limitations of previous ATTR transgenic mouse models. The model demonstrated significantly increased human TTR levels in mouse plasma with Congophilic amyloid deposits in the murine heart and kidneys. Transmission electron microscopy identified both immature and mature amyloid fibrils in the ATTR murine hearts. RNA-sequencing of murine hearts highlighted the activation of inflammatory pathways, notably upregulation of CXCL- and CCL-chemokines, with a significant increase in CCL5 levels observed at the post-translational level among 48 common inflammatory cytokines in mouse plasma. Our findings suggest the CXCL- and CCL-family chemokines play a role in ATTR pathogenesis, providing essential insights into the inflammatory molecular signature of ATTR-CM. This model paves the way for further *in vivo* studies to explore the complex molecular mechanism of ATTR-CM, offering a basis for identifying potential therapeutic targets for ATTR patients.

## Materials and Methods

### AAV Vector Production

Research-grade, high-titer vectors were produced, purified, and titrated by the MEEI/ SERI Gene Transfer Vector Core (https://www.vdb-lab.org/vector-core/). Vector preparations were generated by polyethylenimine or PEI (Polysciences, Cat #24765-2) triple transfection of pACM-Delta ITR-flanked transgene, pKan2/11 (AAV2 rep and AAV11 capsid construct), and pALD-X80 adenoviral helper plasmid in a 1:1:2 ratio, respectively, in HEK293 cells. DNA was transfected in 10-layer HYPERFlasks using a PEI-Max/DNA ratio of 1.375:1 (v/w). 3 days after transfection, vectors were harvested from the HYPERFlasks using Benzonase (EMD Millipore, catalog no. 1016970010) to degrade respective nucleic acids 24 hours after harvesting, the vectors were concentrated by tangential flow filtration and purified by iodixanol gradient ultracentrifugation as previously described. Vaccine candidates were quantified by ddPCR according to a previously published protocol.

### Animal Procedures

All animal experiments were carried out by IACUC at Stanford under animal protocol APLAC 34308. Transgenic TTR-KO mice were ordered through Jackson Laboratories (Strain: B6;129S6-Ttr^tm1Wsb/J^, Stock#: 002382). Humanized TTR V122I mice in a null Ttr background were generated by injection of AAV8-V122I virus (2.5×10^11^ gc/100µL saline) alongside a age- and gender-matched control mice cohort injected with an AAV-GFP (2.5×10^11^ gc/100µL saline). Intravenous injection was administered at 72 weeks (n = 6-12/group) to mimic the age-related onset of symptoms observed in V122I patients. To dynamically monitor the human V122I TTR expression and cardiac function, the echocardiograms and blood draws were taken at baseline, 1 month, 2 months, and 3 months, respectively. At 3 months being our designated endpoint, all mice were sacrificed for tissue harvest alongside extensive histological and transcriptomic analyses.

### Western Blotting

Mouse serum samples were obtained at a monthly time point to monitor plasma TTR expression throughout a 3-month timeframe. Mice serum samples were obtained through retro-orbital blood collection and centrifuged to collect the plasma. Mice plasma protein concentration was quantified by using the Bio-Rad™ DC Protein Assay kit (Reagent A; 5000113, Reagent B; 5000114). 50µg or 2µl plasma samples were prepared with 2x Laemmli buffer (Bio-Rad, Cat # 161-0737) and 2-mercaptoethanol and incubated at 95°C. Proteins were separated by SDS-PAGE and dry transferred to a polyvinylidene difluoride (PVDF) membrane (Immobilon-FL, Cat # IPFL00010, Lot # R7PA8264E). Blocking was performed with an Intercept (TBS) blocking buffer (Thermo, 92760001). To verify the presence of TTR, membranes were incubated overnight at 4°C with rabbit anti-human FLAG (1:1000; Cell Signaling Technology DYKDDDK Tag, 2368S) or Recombinant Anti-Transferrin antibody (Abcam, ab278498) as loading control for mouse plasma. The detection was enabled by using goat anti-rabbit secondary antibody (1:20,000; LICOR C80718-15) and visualized with LICOR ODYSSEY CLx (LICOR) at 84µm with high resolution. Immunoblot quantitative analysis was carried out with the Image Lab 6.1 software (Bio-Rad).

### Human Prealbumin ELISA

The Human Prealbumin ELISA Kit (Abcam, ab231920) is a single-wash 90-minute sandwich ELISA designed for the quantitative measurement of prealbumin protein in serum. Quantitate human prealbumin with 6.3 pg/ml sensitivity. To further validate the concentration of human TTR in mice plasma samples, the Human Prealbumin Enzyme-linked Immunosorbent Assay (ELISA) kit was used (Abcam; ab108895) according to the manufacturer’s protocol. Samples were prepared using a 1:50 ratio for V122I ATTR and a 1:25 ratio for GFP control, and absorbance values were extrapolated from a standard curve by using bovine serum albumin (BSA, Thermo Scientific Catalog B14). Standards were serially diluted to form concentrations of (ng/ul) 31.25, 7.813, 1.953, 0.488, 0.122, and 0. Using a Costar Clear 96 well plate, 50ul of sample/standard were added to each of the wells, and duplicates were made to average the absorbance of each well. Samples were incubated at room temperature before the addition of 1X Biotinylated Prealbumin antibody to each well (1 hour). Washing was repeated after antibody incubation 50ul of 1X Sconjugate was added to each well for a total incubation time of 30 mins at room temperature washing steps were repeated a third time, and 50 ul of chromogen substrate was added to each 20 minutes before terminating the reaction with 50ul of Stop solution. Immediately after observing the blue-to-yellow color change indicating that the reaction had stopped, absorbance at 450 nm was measured with the Spectra Max M5 (Molecular Devices) system and the SoftMax Pro 7.0.3.

### Echocardiography

Left ventricular (LV) functional parameters were assessed by transthoracic echocardiography. Echocardiography was performed at baseline, 1 month, 2 months, and 3 months in both ATTR and control mice using the FUJIFILM Visual Sonics Vevo 3100 (Fujifilm VisualSonics). The cardiac short axis (papillary level), the LV anterior wall end-diastolic thickness (LVAWd), the LV anterior wall end-systolic thickness (LVAWs), the LV end-diastolic internal dimension (LVIDd), the LV end-systolic internal dimension (LVIDs), the LV posterior wall end-diastolic thickness (LVPWd), the LV end-systolic posterior wall thickness (LVPWs), the ejection fraction (EF), and fractional shortening (FS) were measured. The heart rate was also recorded. Echocardiographic measurements were averaged from at least three separate cardiac cycles. Analysis was performed using the Vevo LAB 5.6.1. software in tandem with GraphPad Prism 9.

### Immunohistochemistry

Upon endpoint, mice were euthanized which the heart, lungs, kidneys, spleen, liver, large intestine, tongues, and skeletal muscle were excised and either fixed in 10% formalin, frozen immediately in liquid nitrogen, and stored at −80°C, or conserved in Optimal cutting temperature (OCT) for cryosectioning. The heart was perfused until diastole. Heart weight was recorded and sectioned prior to freezing; the midpapillary level was kept for histological studies and was fixed in 10% formalin for 48 hours before being placed in 70% EtOH in the 4°C fridge. Heart tissues were harvested from adult mice and immediately fixed in formaldehyde to preserve structural integrity. Following fixation, tissues were embedded in paraffin and sectioned at 4 µm. The sections were deparaffinized in the following procedure: xylene wash twice for 4 minutes each time, 100% ethanol (EtOH) twice for 1 minute each time, 95% EtOH for one minute, 90% EtOH for one minute, and ddH20 three times and one minute each time. Samples were deparaffinized and rehydrated by a descendant ethanol series and incubated with 1% NaOH in 80% EtOH three times.

### Masson’s Trichrome Staining

Murine hearts were excised and fixed in 10% formalin overnight, followed by 4 µm thick sections embedded in paraffin mounted on glass slides. After deparaffinization and rehydration, the murine cardiac tissue was stained with Weigert’s iron hematoxylin, Biebrich scarlet acid fuchsin, and aniline blue from the Masson’s Trichrome Staining Kit (Polysciences, 25088). Samples were visualized at 20x magnification under Brightfield with the Zeiss LSM800 Airyscan (Zeiss) with 5-10 separate views per sample. Fibrosis and amyloid fractional percentages were measured with ImageJ.

### Thioflavin T Staining

The sections were deparaffinized and hydrated through a series of graded alcohol washes, followed by incubation with 0.5% of thioflavin T (Thermo Scientific, Cat # 211760050, Lot # A0451794) in 0.1 N HCl. In brief, the sections are deparaffinized and rehydrated. Using a hydrophobic marker, the specimen section and an adjacent positive control section are encircled on the top of the slides. A drop of the working solution of thioflavin T is placed on the slides, which are subsequently kept in a humidity chamber for 15 min. The slides are rinsed in deionized water (for 5 min) and kept in deionized water while cover slipping with Aqua mount. The frozen sections are rinsed two times in PBS (for 10 min each) before incubation with thioflavin T and, thereafter, rinsed sequentially in PBS and deionized water (for 5 min each) and cover slipped. Thioflavin stain is viewed under a fluorescein gate using a standard immunofluorescence microscope. Thioflavin T fluoresces only when bound to amyloid fibrils, where it shows a bright yellow-green fluorescence. In contrast, in the absence of amyloid deposits, the dye fluoresces faintly.

### Congo Red Staining

The detection of amyloid deposits in 4µm sections of mouse cardiac tissue was verified by Congo red (Sigma, Cat # C6277-25G, Lot # MKCL0814) staining under brightfield and polarized light, respectively. Slides were then stained with hematoxylin, rinsed, and finally stained with 0.2% Congo red in 1% NaOH in 80% EtOH. Samples were visualized with the Zeiss LSM800 Airyscan (Zeiss).

### Transmission electron microscopy (TEM)

Ultrastructural analysis of the samples was conducted at Stanford Cell Sciences Imaging Facility using the (JEOL, JEM-1400) transmission electron microscope (TEM) operated at an accelerating voltage of 120 kV. The specimens were first fixed with 2.5% glutaraldehyde, post-fixed in osmium tetroxide, dehydrated in a graded series of ethanol solutions, and embedded in epoxy resin. Ultrathin sections (approximately 70 nm) were cut with a diamond knife, collected onto copper grids, and stained with uranyl acetate and lead citrate to enhance contrast. The grids were then examined under the TEM, and images were captured using a CCD camera. This system allowed for the high-resolution examination of the cellular ultrastructure, providing insights into the morphological changes at the nanometer scale at 10000x, 25000x, and 40000x, respectively.

### RNA extraction and cDNA synthesis

The total RNA of the cells was isolated using the Trizol RNA extraction method. In brief, 100 mg of mouse liver from ATTR and control mice were mixed with 1 mL TRIzol™ Reagent (Thermo Fisher Scientific, Cat# 15596018) and then homogenized using a Tissue Lyser II (Qiagen) set at 2 minutes, 20 Hz. until no precipitate could be observed. The tissue lysate was then mixed with 200 μL of chloroform and centrifuge 12,000 × g at 4 °C for 15 min. The aqueous phase was collected, mixed with 500 μL of isopropanol, and incubated at −20 °C for 30 min to precipitate RNA. The RNA pellet was harvested by centrifugation at 4 °C and 15,000 g for 30 min, washed with 70% EtOH, air-dried, and resuspended in 50 μL RNase-free water. For cDNA synthesis, 500 ng of RNA was taken using iScript RT Supermix (BioRad; Cat# 1708840) for qPCR.

### Real-time quantitative polymerase chain reaction (RT-qPCR)

RT-qPCR was conducted to quantify humanized TTR expression in mice liver tissues at a 3-month endpoint. For quantitative analysis, the PCR reaction was carried out with an initial step of the following thermal cycling profile: 95 °C for 20 sec, followed by 40 cycles of amplification (95 °C for 10 s, 60 °C for 20 s, 70 °C for 10 s) on the CFX96 Touch Real-Time PCR Detection System (BioRad, 1845096) using the CWBIO UltraSYBR Mixture (Cat.CW0957L). Specific primers for human V122I TTR and GAPDH (housekeeping gene) were listed in Supplementary Information **(Sup Fig 3A)**. A melting curve analysis was conducted to validate the specificity of RT-qPCR products. The relative CT value was determined and normalized against the expression level of endogenous human glyceraldehyde 3-phosphate dehydrogenase (GAPDH). The expression quantities of the above genes were rectified with GAPDH as relative quantities by 2−ΔΔCT methods.

### RNA Library Construction, Quality Control and Sequencing

RNA extraction from murine hearts was performed using a miRNeasy Mini Kit (Qiagen, 217004). RNA quantification was performed using the Thermofisher Scientific Nano-drop One machine. All the procedures were carried out following the specifications of the manufacturer. Library Construction: Messenger RNA was purified from total RNA using poly-T oligo-attached magnetic beads. After fragmentation, the first strand of cDNA was synthesized using random hexamer primers, followed by the second strand of cDNA synthesis using either dUTP for directional library or dTTP for non-directional library [41]. For the non-directional library, it was ready after end repair, A-tailing, adapter ligation, size selection, amplification, and purification. The library was checked with Qubit and real-time PCR for quantification and bioanalyzer for size distribution detection. Quantified libraries will be pooled and sequenced on Illumina platforms, according to effective library concentration and data amount. RNA sequencing and downstream analysis were performed by Stanford Genome Center.

### Transcriptomic Analysis

#### Data Quality Control

Quality control of raw sequencing data was performed using FastQC, which provided initial read quality assessment. Subsequently, MultiQC was used to aggregate the FastQC reports into a single comprehensive document. This step allowed for the evaluation of key quality metrics such as per base sequence quality, per sequence quality scores, per base N content, sequence length distribution, and sequence duplication levels. Based on these quality assessments, sequences were trimmed and filtered to remove adapters and low-quality bases using Trimmomatic.

#### Read Alignment and Quantification

Sequencing reads were aligned to the mouse reference genome (specify version, GRCm39) using the Spliced Transcripts Alignment to a Reference (STAR) aligner. STAR was configured to employ default parameters for aligning RNA-seq data, ensuring accurate alignment around splice junctions. Following alignment, the featureCounts tool from the Subread package was utilized to quantify gene-level read counts, summarizing reads per gene based on their overlap with annotated exons.

#### Data Preprocessing and Normalization

Raw read counts were loaded into R using data.table and dplyr packages for preprocessing. Gene names were assigned based on Ensembl IDs extracted from the first file in the “GeneCounts” directory. A count matrix was constructed with samples as columns and genes as rows. Low-expression genes were filtered out using the filterByExpr function from the edgeR package, which applies default thresholds tailored to the library sizes and read depth of the experiment. Normalization factors were calculated to adjust for library size differences among samples using the TMM (Trimmed Mean of M-values) method implemented in the edgeR package. The resulting normalized counts were used for downstream differential expression analysis.

#### Differential expression analysis

The design matrix was constructed to model the effects of treatment groups. Using the voom function from the limma package, read counts were transformed to log-counts per million (log-CPM), stabilizing variance across the range of counts. The linear models for microarray data (limma) framework were applied, using empirical Bayes moderation to improve the estimation of model coefficients and their variances. Contrasts were set up to test for specific comparisons of interest (e.g., ATTR vs. control).

#### Pathway Analysis

DEGs were subjected to pathway and gene ontology analysis using the enrichR and pathfindR packages to identify enriched biological themes and pathways. Results were visualized using enrichment charts, and clustered to identify patterns among significantly enriched terms.

#### Visualization

EnhancedVolcano plots were used to visually represent differentially expressed genes (DEGs), highlighting genes with significant fold-changes and adjusted p-values. An interactive multidimensional scaling (MDS) plot and heatmaps of DEGs were also created to explore sample relationships and expression patterns, respectively.

### Mouse Plasma Immunoassay

Luminex Thermo-Fisher/Life Technologies Mouse 48 plex kits were used for immunoassay at Stanford Human Immune Monitoring Center (HIMC). This assay was performed by the Human Immune Monitoring Center at Stanford University. Mouse 48 plex Procarta kits (EPX480-20834-901) were purchased from Thermo-Fisher/Life Technologies, Santa Clara, California, USA, and used according to the manufacturer’s recommendations with modifications as described. Briefly, beads were added to a 96-well plate and washed in a Biotech ELx405 washer. Samples were added to the plate containing the mixed antibody-linked beads and incubated overnight at 4°C with shaking. Cold (4 °C) and room-temperature incubation steps were performed on an orbital shaker at 500–600 rpm. Following the overnight incubation, plates were washed in a BioTek ELx405 washer and a biotinylated detection antibody was added for 60 minutes at room temperature with shaking. The plate was washed as described and streptavidin-PE was added for 30 minutes at room temperature. The plate was washed as above and reading buffer was added to the wells. Each sample was measured in duplicate or singlet. Plates were read using a Luminex 200 or an FM3D FlexMap instrument with a lower bound of 50 beads per sample per cytokine. Custom Assay Chex control beads were purchased from Radix BioSolutions, Georgetown, Texas; and added to all wells. The average median fluorescence intensity (MFI) was utilized for the quantitative analysis.

### Statistics

Statistical analysis was performed by GraphPad Prism 9.4.0. (Graph Pad Software, San Diego, CA, USA). In each experiment, all determinations were performed at least in duplicates. Two-tailed unpaired Student’s t-tests were used to evaluate the statistical significance of differences between the two groups. The analysis of variance (ANOVA) test was applied for the comparison between more than two groups. The data were presented in the format of mean ± standard error of the mean (SEM). Statistical significance was considered at P < 0.05. Statistical parameters and corresponding P values are included in the figure legends.

## Results

### Humanized ATTR Mouse Model Generation

To elucidate the role of inflammation in ATTR, we first established and validated an *in vivo*, humanized TTR V122I mouse model of ATTR. We utilized a murine *TTR* knockout (TTR-KO) mouse (Jackson Laboratories, Stock#:002382) to overcome the drawback and enhance the expression of human *TTR* in our mouse model (6-12 mice/ group for ATTR and placebo). Next, to better mimic the pathophysiology of human TTR production in the liver, recombinant Adeno-associated Virus (AAV) serotype 8 (2.5×10^11^ gc/100µL saline) was utilized as human *V122I* gene delivery vectors to specifically target the liver of TTR-KO mouse, which can transduce over 90% of hepatocytes in the mouse liver following intravenous injection [42]. As V122I ATTR was shown to predominantly affect elderly males [43], to study the gender difference both male and female mice were included in the study. As ATTR is an aging-related disorder, the experimental design of our *in vivo* mouse model is based on prior clinical research supporting the claim that amyloid fibrils are most clearly observed in humans after the age of 65 years old [1, 44]. In brief, aged TTR-KO male and female mice were randomly divided into two subgroups at 72 weeks of age. The ATTR group received AAV8-V122I from Gene Transfer Vector Core (GTVC) at Harvard Medical School (2.5×10^11^ gc/100ul saline), while the age-, sex-matched control group received AAV8-GFP from Addgene (2.5×10^11^ gc/100ul saline). Cardiac function, circulating TTR expression level, and amyloid deposition were determined longitudinally for three months post-injection.

### Elevated Human TTR Expression in Male and Female ATTR Mice Post Injection

To dynamically track circulating human *TTR* expression level during the time course. Mouse plasma was monthly collected by retro-orbital approach, western blot, and human TTR ELISA was performed to assess the circulating TTR level. The western blots of the mouse plasma from male ATTR, and female ATTR mice showed significantly increased serum human TTR expression during the time course post-injection 1-,2-, and 3 months compared to the control mice **(Fig 1A-B) (Sup Fig 2A)**. Next, to measure human TTR concentration in the plasma more precisely, we also performed human TTR ELISA. The quantitative analysis of human TTR ELISA in male and female mice showed dramatically elevated human TTR levels in the ATTR group compared with their controls (human TTR level (ng/ml) Male ATTR: 109.9 ± 5.568; Male control: 28.17 ± 7.010; p=0.0008, N=3 mice/group; Female ATTR: 127.5 ± 32.43; Female control: 20.08 ± 8.351; p=0.0327, N=3 mice/group) **(Fig 1C-D)**, which validated the significant increased TTR level in both male and female ATTR mice post-injection. When we compared TTR expression level between male and female ATTR mice, there was no significant difference post-injection at 3 months **(Fig 1E)**. We also performed the RT-qPCR to monitor the human TTR expression efficacy, we did not observe any significant difference in human TTR expression in the liver between male and female ATTR mice which was consistent with our ELISA findings (Sup Fig 3B).

**Figure 1.**
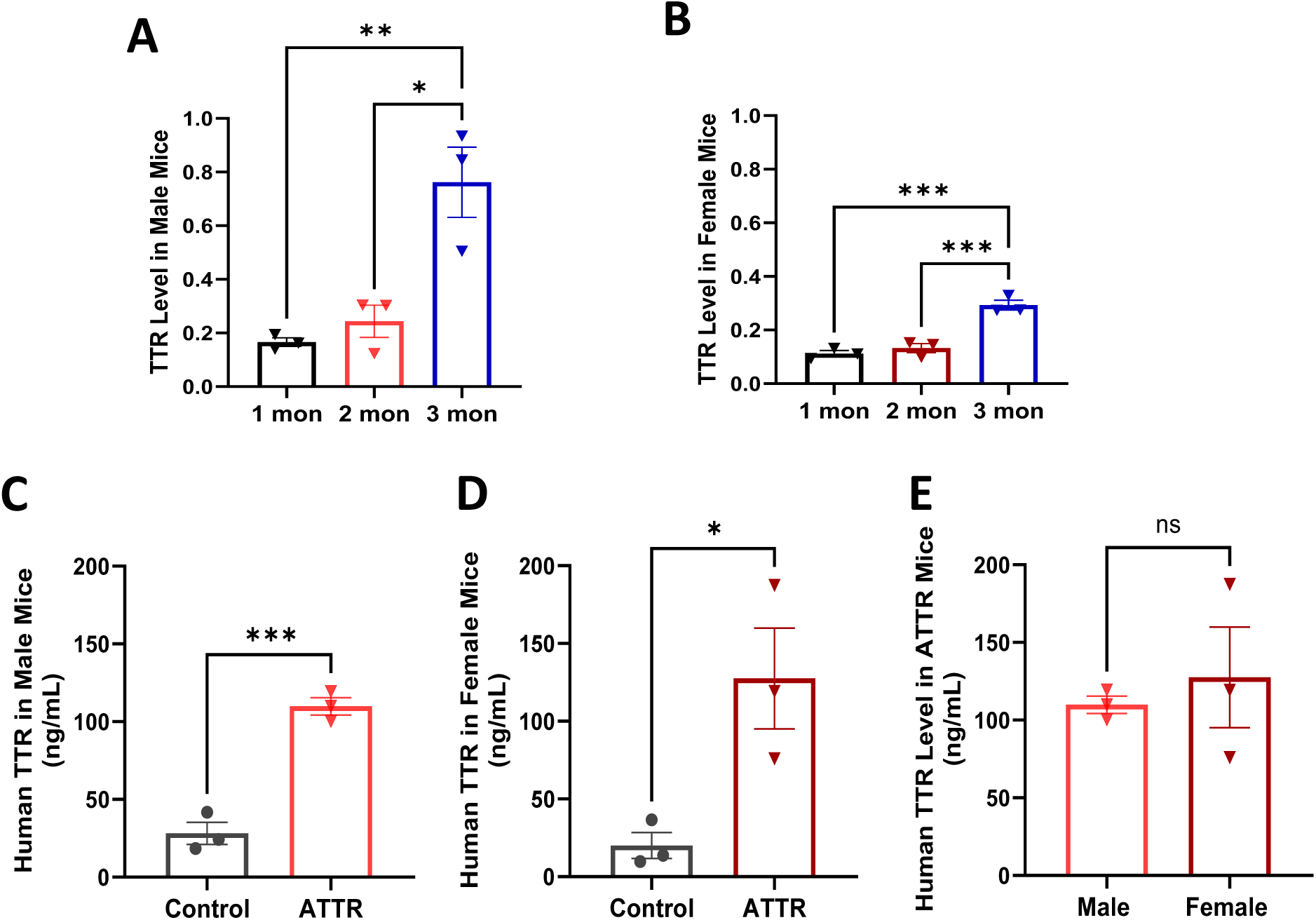
Elevated Human TTR Expression in Male and Female ATTR Mice Post Injection. **(A-B)** Quantitative analysis of western blots of the mouse plasma from male ATTR, female ATTR mice, and control mice during time course post-injection 1-,2-, and three months showed significantly increased serum human TTR expression. Transferrin was used as a loading control for the mouse plasma western blots. N=3 mice/group; Data are presented as mean ± SEM. *p < 0.05, **P < 0.01, ***P < 0.001 by two-way ANOVA. **(C-D)** Quantitative analysis of human *TTR* ELISA in male and female mice showed dramatically elevated TTR levels in the ATTR group compared with controls. N=3 mice/group; Data are presented as mean ± SEM. *p < 0.05, ***P < 0.001 by nonparametric T-test. **(E)** In a human TTR expression level comparison in male and female ATTR mice, there was no significant difference in human TTR level between male and female ATTR mice after three months of injection. N=3 mice/group; Data are presented as mean ± SEM by nonparametric T-test.

### Preserved Cardiac Function and Structure in Male and Female ATTR Mice Post Injection 3 months

At 3 months post-injection, we did not observe any significant changes in cardiac structure or function **(Fig2, supp Fig 1G-H).** There was a preserved cardiac function in male and female ATTR mice (FS% Male ATTR: 26.07 ± 3.667; Male control: 22.69 ± 1.585; p=0.3712, N=6-8 mice/group; Female ATTR: 26.62 ± 1.980; Female control: 31.25 ± 4.482; p=0.3397, N=5-6 mice/group) compared with controls post-injection 3 months. In addition, we harvested murine organs for downstream histology studies. The heart-to-body weight (HW/BW) coefficient has been used for characterizing myocardial hypertrophy [45]. The lung-to-body weight ratio (LW/BW) was commonly utilized to validate the pulmonary hypoplasia [46]. Our data showed that the cardiac structure was maintained in the male and female ATTR mouse which we did not observe cardiac hypertrophy (HW/BW Male ATTR:6.139 ± 0.1168, Male Control:5.351 ± 0.3235, p=0.1027, n=3-4 mice/group; Female ATTR:5.595 ± 0.8755, Female Control:4.331 ± 0.6241, p=0.3049, n=3-4 mice/group) **(Fig 3A-B)** nor pulmonary edema (LW/BW Male ATTR:5.783 ± 0.3484, Male Control:4.603 ± 0.4303, p=0.1004, n=3-4 mice/group; Female ATTR:6.504 ± 0.5724, Female Control:4.963 ± 0.8239, p=0.1993, n=3-4 mice/group) **(Fig 3E-F)** in ATTR mice compared with control. We further validated our measurements by using heart weight to tibial length (HW/TL) **(Fig 3C-D)** and lung weight to tibial length (LW/TL) **(Fig 3G-H)**, which was consistent with the previous findings. In addition, we also monitored the morphology **(Sup Fig 1A-D)** and their body weight changes **(Sup Fig 1E-F).** Altogether, the results suggested preserved cardiac function and maintained cardiac structure in ATTR mouse post-injection 3 months, which may need a longer follow-up study to see if the amyloid fibrils cause detectable cardiac function and structural changes.

**Figure 2.**
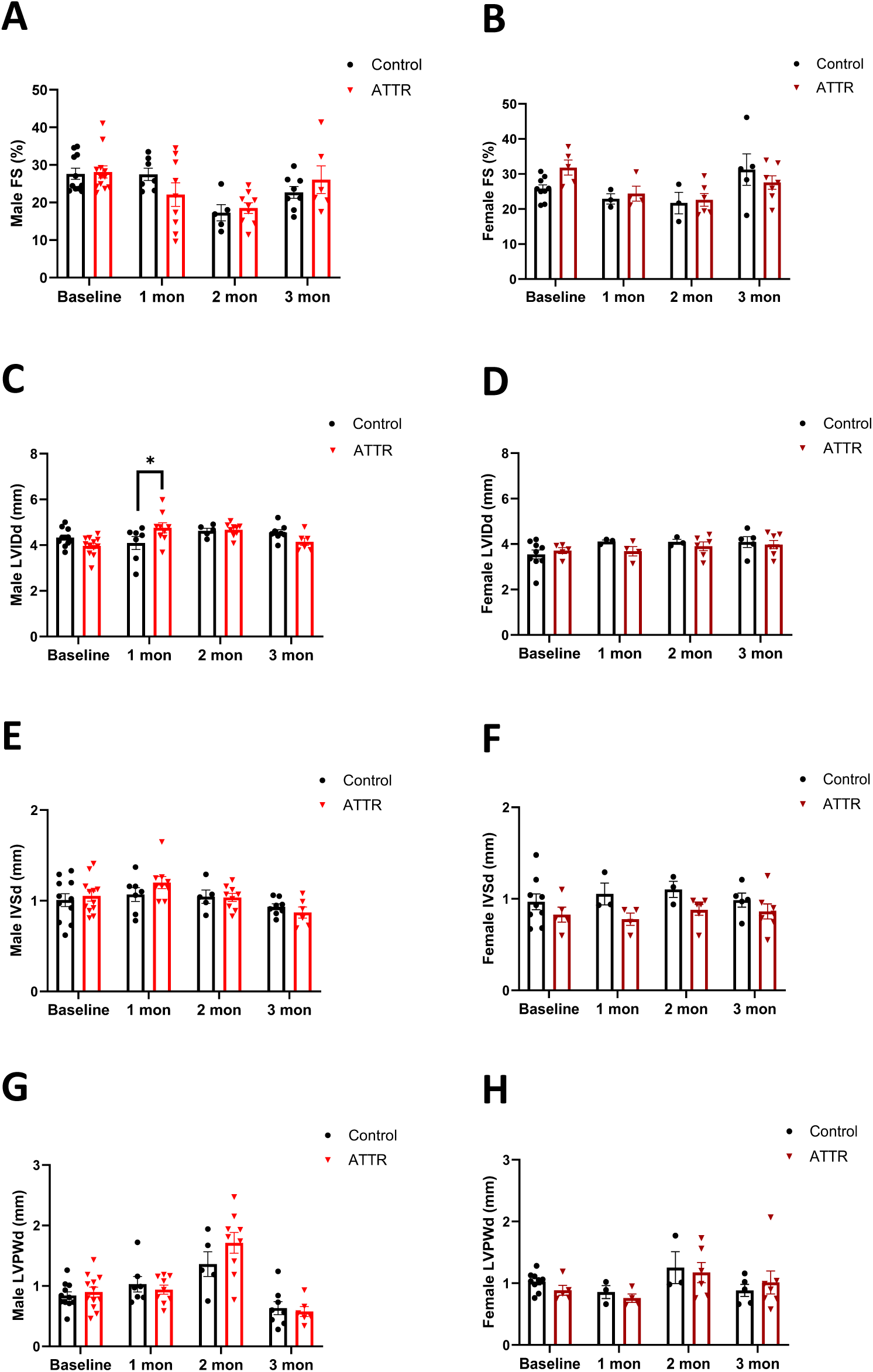
Preserved Cardiac Function in Male and Female ATTR Mice Post Injection 3 months. **(A-B)** Quantitative analysis of fractional shortening (FS%) in male and female mice post injection during time course 3 months; **(C-D)** Quantitative analysis of left ventricular internal diameter end diastole (LVIDd) in male and female mice post injection during time course 3 months**; (E-F)** Quantitative analysis of interventricular septum thickness end diastole (IVSd); **(G-H)** Quantitative analysis of left ventricular posterior wall thickness end diastole (LVPWd) in male and female mice post injection during time course 3 months. The quantitative analysis of echocardiograms showed no significant changes for FS%, LVIDd, IVSd, and LVPWd in male ATTR and female ATTR compared to their controls. N=3-12 mice/group; Data are presented as mean ± SEM. *p < 0.05 by two-way ANOVA.

**Figure 3.**
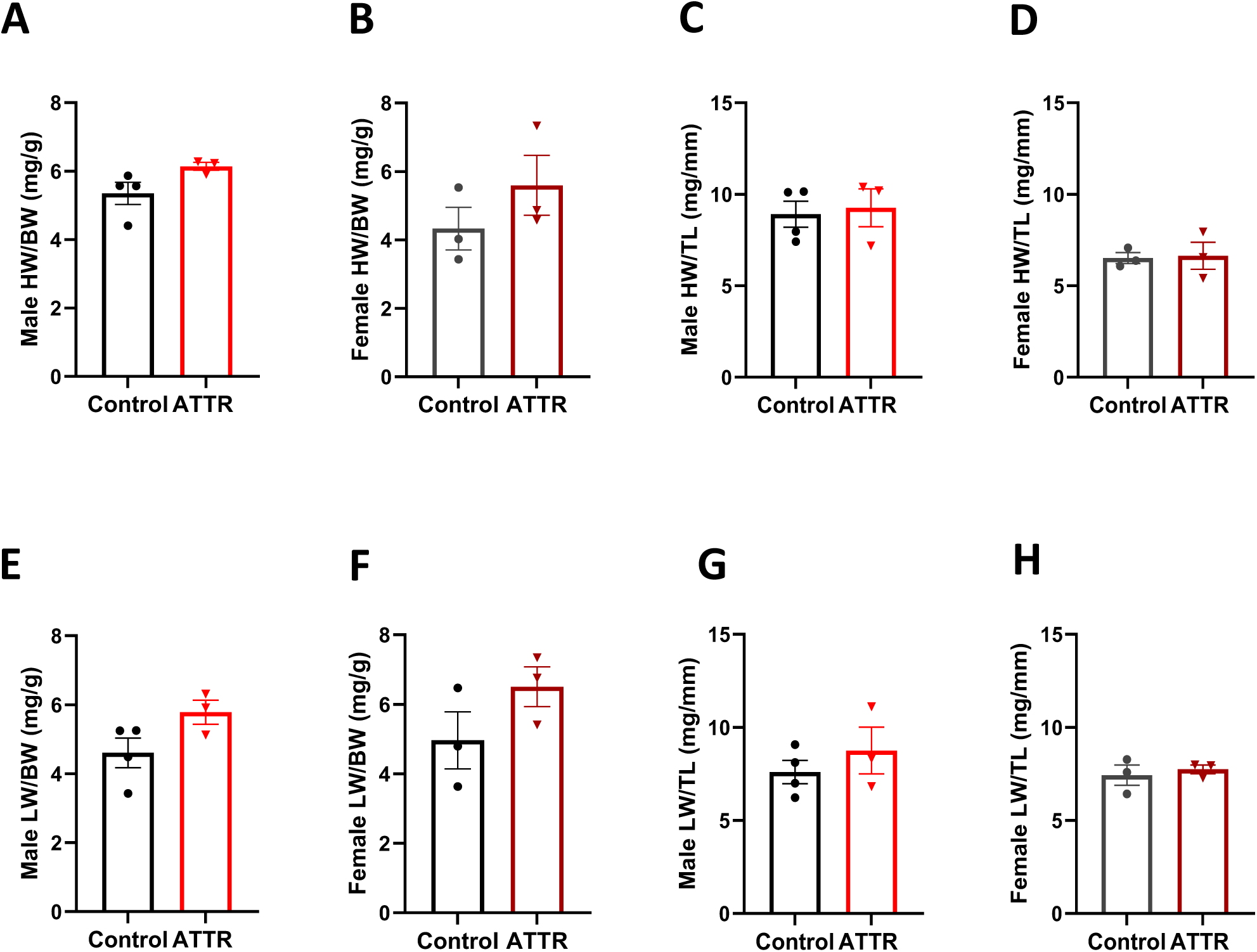
Maintained Cardiac Structure in Male and Female ATTR Mice Post-injection 3 months. **(A-B)** Quantitative analysis of mouse heart weight to body weight ratio (HW/BW) from male and female ATTR mice compared with controls; **(C-D)** Quantitative analysis of mouse heart weight to tibial length (HW/TL) ratio from male and female ATTR mice compared with controls; **(E-F)** Quantitative analysis of mouse lung weight to body weight ratio (LW/BW) from male and female ATTR mice compared with controls; **(G-H)** Quantitative analysis of mouse lung weight to tibial length (LW/TL) ratio from male and female ATTR mice compared with controls. Statistical analysis of organ measurement shows no significant changes in HW/BW ratio, HW/TL ratio, LW/BW ratio, and LW/TL in males and females in the ATTR group compared with controls. N=3-4 mice/group; Data are presented as mean ± SEM.

### Cardiac Amyloid Deposition in ATTR Mouse Hearts

Next, to detect the cardiac amyloid deposition, we performed multiple histology studies. By using Masson’s trichrome staining, the connective fibrous tissue is evidenced by intense blue staining, whereas amyloid deposits show a blue-violet color **(Fig 4A-C),** which was consistent with the previous report. The quantitative images analysis showed a significantly increased fibrosis and amyloid deposition in ATTR murine hearts (Extent of amyloid and fibrosis % of biopsy tissue ATTR: 0.4655 ± 0.0824; Control: 0.2922 ± 0.0308; p<0.05, n=3 mice/group) **(Fig 4F).** In addition, we performed two amyloid-specific stains to further validate the cardiac amyloid deposition in the ATTR mice. Congo red staining is commonly used to detect amyloid fibrils, which present red under bright light, while shown as apple-green birefringence under polarized light, which is due to the β-pleated sheet structure of the fibrils [47–51]. By using Congo Red staining, we observed the modest cardiac amyloid deposition in the ATTR mouse **(Fig 5B, E)** compared with the human ATTR-CM hearts, which served as a positive control **(Fig 5A, D)** and their murine controls hearts **(Fig 5C, F)**. Furthermore, we also detected the renal amyloid deposition in ATTR mouse **(Sup Fig 4A, C)**, which indicated a systemic amyloid deposition in the ATTR mouse model. Lastly, a more sensitive Thioflavin T staining (ThT) was also employed in murine hearts as described [52], which further validated the cardiac amyloid deposition **(Sup Fig 5B-D)** in ATTR mouse.

**Figure 4.**
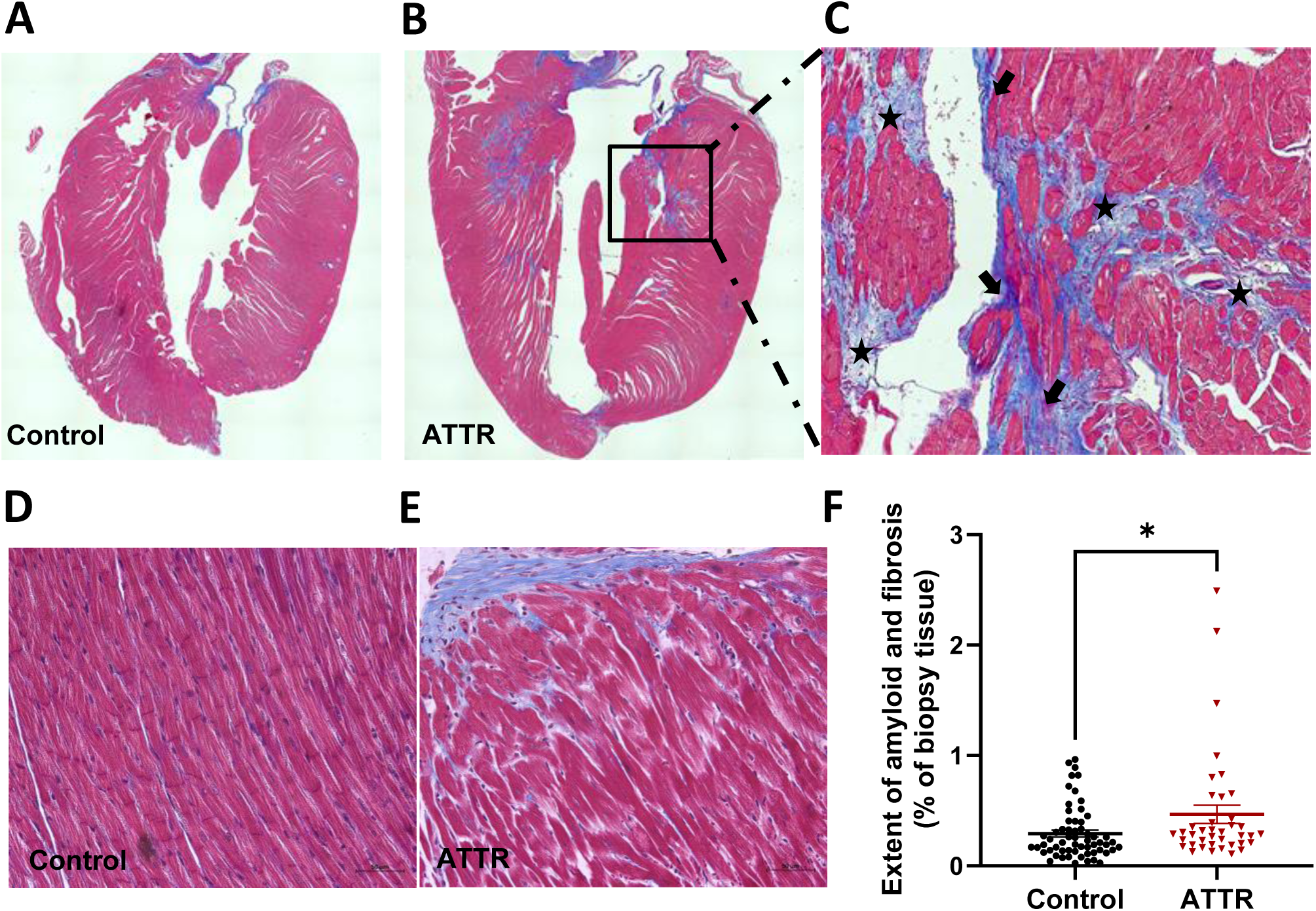
Increased fibrosis and Amyloid by Masson’s Trichrome Staining. **(A-E)** Representative Masson’s Trichrome staining (MTS) findings on endomyocardial biopsy of whole hearts and partial hearts (slides digitized at 20x) from male ATTR and control mice. **(C)** Masson’s trichrome staining shows that the connective fibrous tissue (arrows) is evidenced by intense blue staining, whereas amyloid deposits show a blue-violet color (asterisks). Original magnification, 40x. **(F)** Quantitative analysis of MTS for fibrosis and amyloid fibrils showed a significantly increased extent of amyloid and fibrosis in male ATTR hearts compared with controls. Each dot represents one field acquisition (magnification of 20x with a confocal microscope), 60 fields for 3 controls, and 39 for 3 ATTR mice. Data are presented as mean ± SEM. *p < 0.05 by nonparametric T-test.

**Figure 5.**
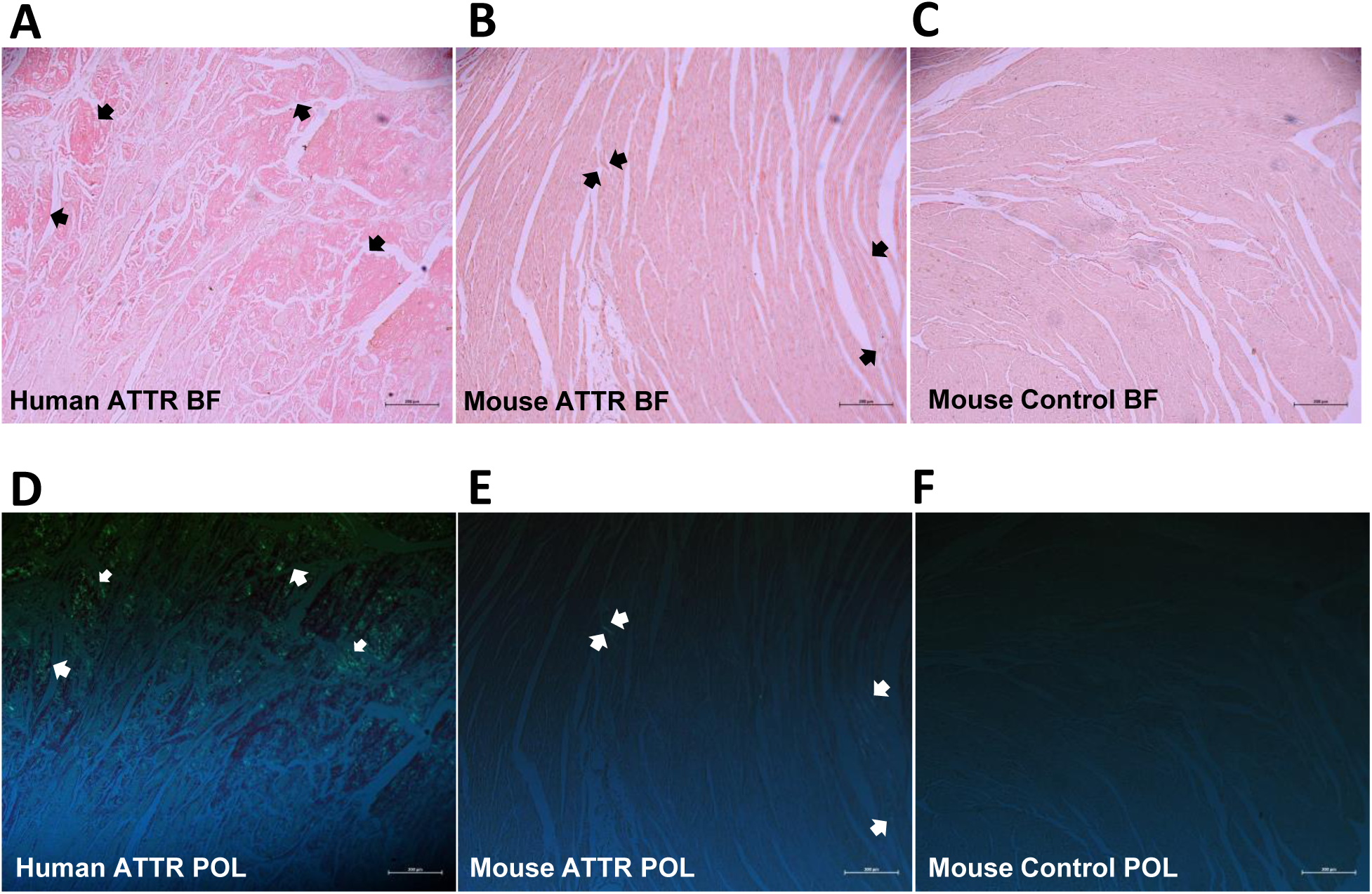
Cardiac Amyloid Deposition in ATTR Mouse Hearts by Congo Red Staining. **(A-C)** Representative images of Congo Red staining for human ATTR heart biopsy, male ATTR mouse, and control mouse hearts under bright field (BF) Original magnification, 5x. The black arrows indicated the amyloid fibrils were stained as red. **(D-F)** Representative images of Congo Red staining for human ATTR heart biopsy, male ATTR mouse, and control mouse hearts under polarized light (POL) imaging. The white arrows indicated the amyloid fibrils were shown as apple green under polarized view. Original magnification, 5x.

### Mature and Immature Amyloid Fibrils in Extracellular Matrix of the ATTR Mouse Hearts Post Injection 3 Months

TEM was employed to confirm the ultrastructure of amyloid fibrils **(Fig 6)**. The previous ultrastructural study of amyloid fibrils using biopsy or autopsy specimens from hereditary transthyretin amyloidosis patients indicated that amyloid fibrils seemed to mature from dotty structures among amorphous materials in the peripheral nervous system [53]. TEM of the murine hearts at the high-powered view (10,000x,25,000x,40,000x) revealed both mature and immature amyloid fibrils in ATTR hearts compared with its controls. We observed the immature fibrils were shown as dotted structures while the mature fibrils were long, thin, mesh-like structures in the cardiac extracellular matrix (ECM) of ATTR mouse compared with controls **(Fig 6B-C)**. These results are similar to the previously reported findings.

**Figure 6.**
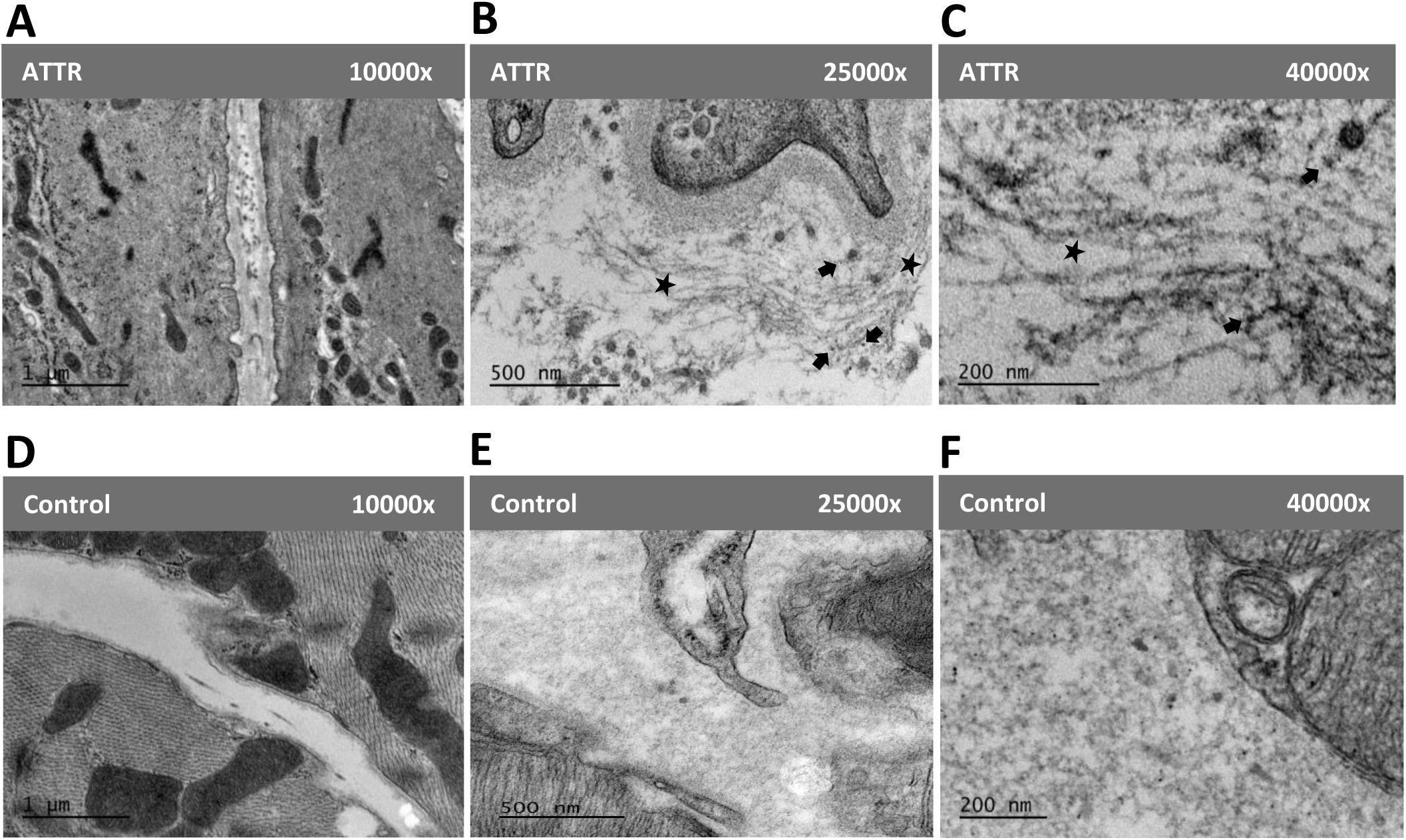
Mature and Immature Amyloid Fibrils in Extracellular Matrix of the ATTR Mouse Hearts. **(A-F**) Representative Transmission electron microscopy (TEM) images for ATTR mouse and control mouse hearts at different magnifications (10,000x,25,000x,40,000x). **(B-C)** TEM revealed both mature (asterisks) and immature amyloid fibrils (arrows) in male ATTR murine hearts; the arrows indicated immature fibrils were shown as dotted structures and mature fibrils were long, thin, mesh-like structures in the cardiac extracellular matrix of ATTR mouse hearts compared with controls.

### Distinct Transcriptomic Patterns in ATTR Mouse Compared with Control

At the endpoint, we collected the aged ATTR mouse hearts and extracted RNA for downstream bulk RNA sequencing at Stanford Genomic Core. The Principal Component Analysis **(Fig 7A)** and the heatmap of the RNA-sequencing analysis **(Fig 7B)** for the ATTR murine hearts and control hearts identified distinct transcriptomic patterns in the ATTR group compared to age- and gender-matched controls. The pathway analysis indicates the upregulation of inflammation signals in male ATTR murine hearts compared with controls, including leukocyte transendothelial migration, T-cell signaling, TNF signaling, and B-cell receptor signaling pathway **(Fig 7C)**. The results suggest that there were distinct transcriptomic patterns in ATTR mice compared with controls.

**Figure 7.**
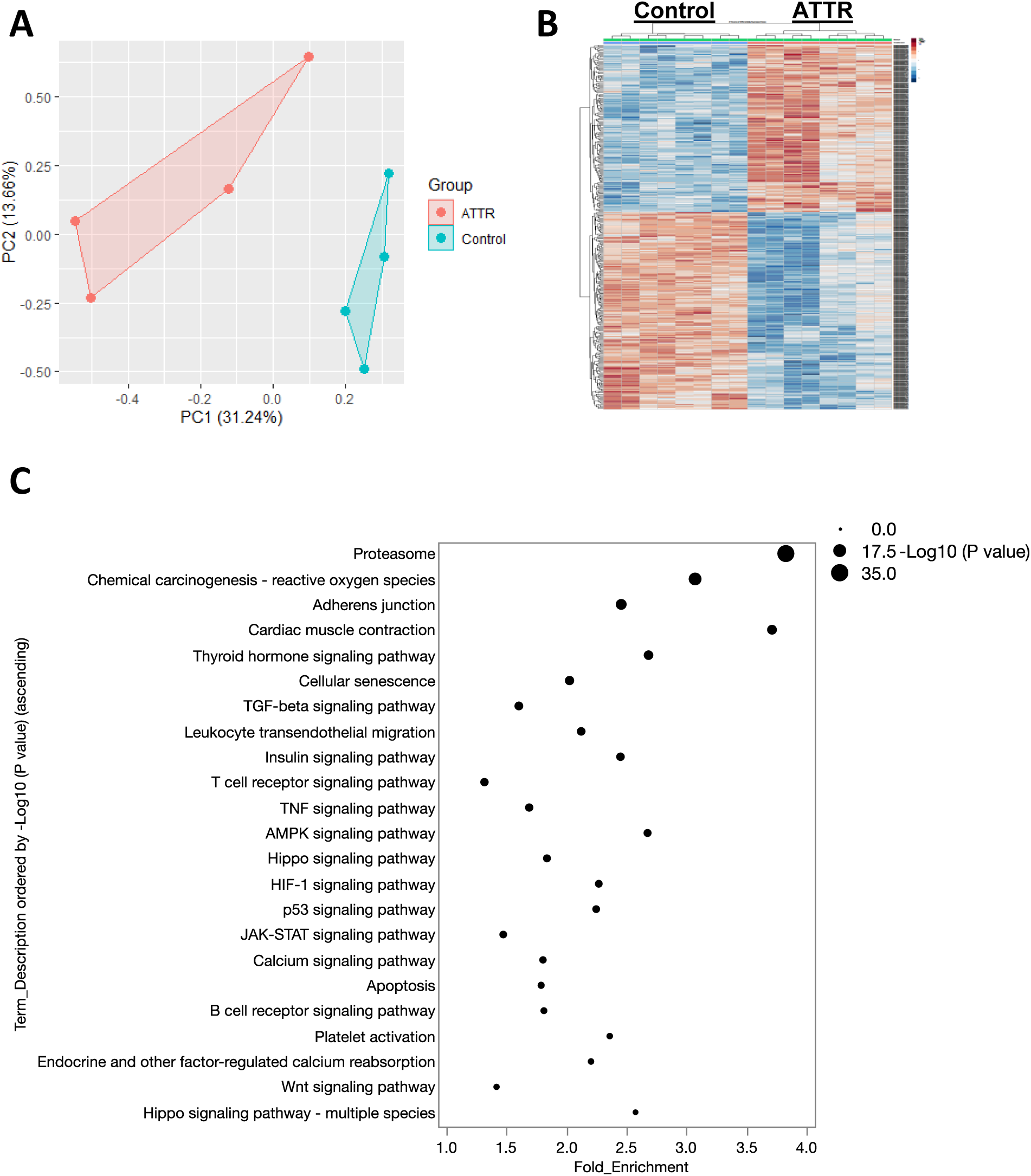
Distinct Transcriptomic Patterns in ATTR Mouse Compared with Control. **(A)** The Principal Component Analysis of male ATTR and control mice; **(B)** The heatmap of the RNA-sequencing analysis for the ATTR murine hearts and control hearts identified distinct transcriptomic patterns in the ATTR group compared to gender-matched controls; **(C)** The pathway analysis indicate upregulation of inflammation signals in male ATTR murine hearts compared with controls, including leukocyte transendothelial migration, T-cell signaling, TNF signaling and B-cell receptor signaling pathway. The pathways (Y axis) were ordered by ascending –Log_10_ (P-values), and the fold enrichment was listed in the ascending X axis.

### Up-regulation of Inflammatory Pathways and Markers in ATTR Murine Hearts

Next, further analysis indicates the upregulation of inflammatory signaling pathways in ATTR murine hearts compared with controls, such as MAPK signaling pathway, TGF-beta signaling pathway, AMPK signaling pathway, HIF-1 signaling pathway, and JAK-STAT signaling pathway **(Fig 8A)**. The differentially expressed genes (DEGs) analysis identified up-regulation of inflammatory markers in ATTR hearts, including CXCL1, CXCL2, CXCL3, and CCL20 **(Fig 8B)**. The role of CCL and CXCL chemokine families in the pathogenesis of cardiovascular disease, particularly heart failure and cardiac amyloidosis, is complex and multifaceted. These chemokines play crucial roles in mediating inflammation, fibrosis, and immune cell recruitment, which are key processes in the development and progression of these conditions [54]. Our data suggest that there was upregulation of inflammation in ATTR mouse heart compared with controls which we also revealed CXCL- and CCL-cytokine family may potentially contribute to the pathogenesis of ATTR.

**Figure 8.**
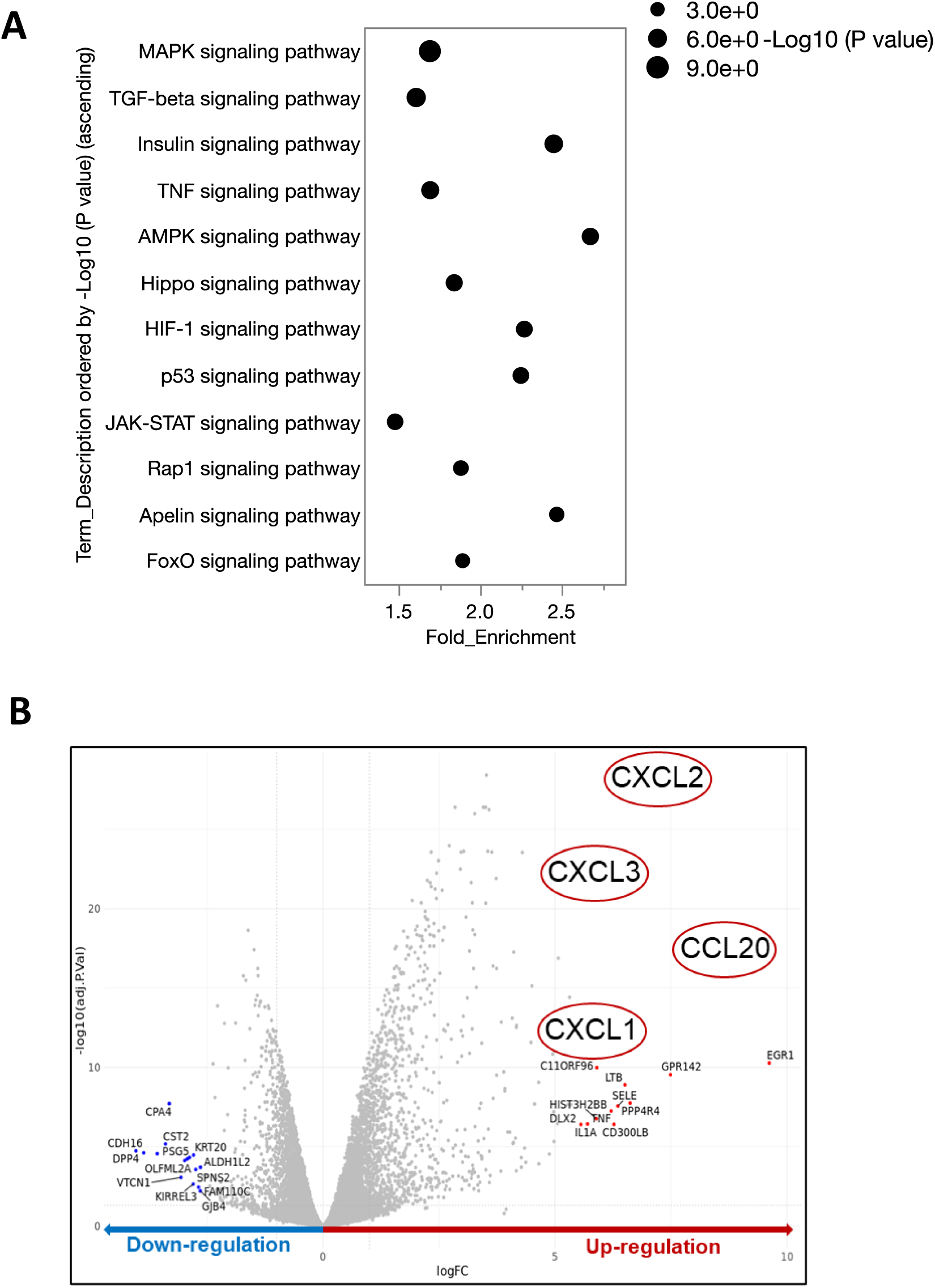
Up-regulation of Inflammatory Pathways and Markers in ATTR Murine Hearts. **(A)** The pathway analysis indicates the upregulation of inflammatory signaling pathways in ATTR murine hearts compared with controls. The pathways (Y axis) were ordered ascending –Log_10_ (P-values), and the fold enrichment was listed in the X axis ascending. **(B)** The differentially expressed genes (DEGs) analysis showed up-regulation of inflammatory markers in ATTR hearts, including CXCL1, CXCL2, CXCL3, and CCL20.

### Elevated Circulating CCL5 in ATTR Mouse Post Injection 3 Months

To dive deeper into the circulating level of CCL and the CXCL cytokines in the ATTR mice, we performed a mouse 48 Plex (Procarta) luminance-based immunoassay to check 48 common inflammatory cytokines in mouse plasma **(Fig 9A)**. The averaged median fluorescence intensity (MFI) across duplicates and four control beads (Radix BioSolutions, Chex1-Chex4) was utilized for quantitative analysis of the assay. We found the tendency of increased CXCL-1 and CCL3 in ATTR mice **(Sup Fig 6)**. Specifically, we observed a significantly increased level of CCL5 (Chemokine C-C motif ligand 5) (MFI ATTR: 801 ± 105; Control: 426± 64; p=0.0061, N=3 mice/group) in ATTR murine plasma compared with controls **(Fig 9B)**. Altogether, our data suggested that there was a potential link between inflammation and ATTR, in which the CXCL and CCL cytokine family may play a pivotal role in mediating the ATTR-derived inflammation.

**Figure 9.**
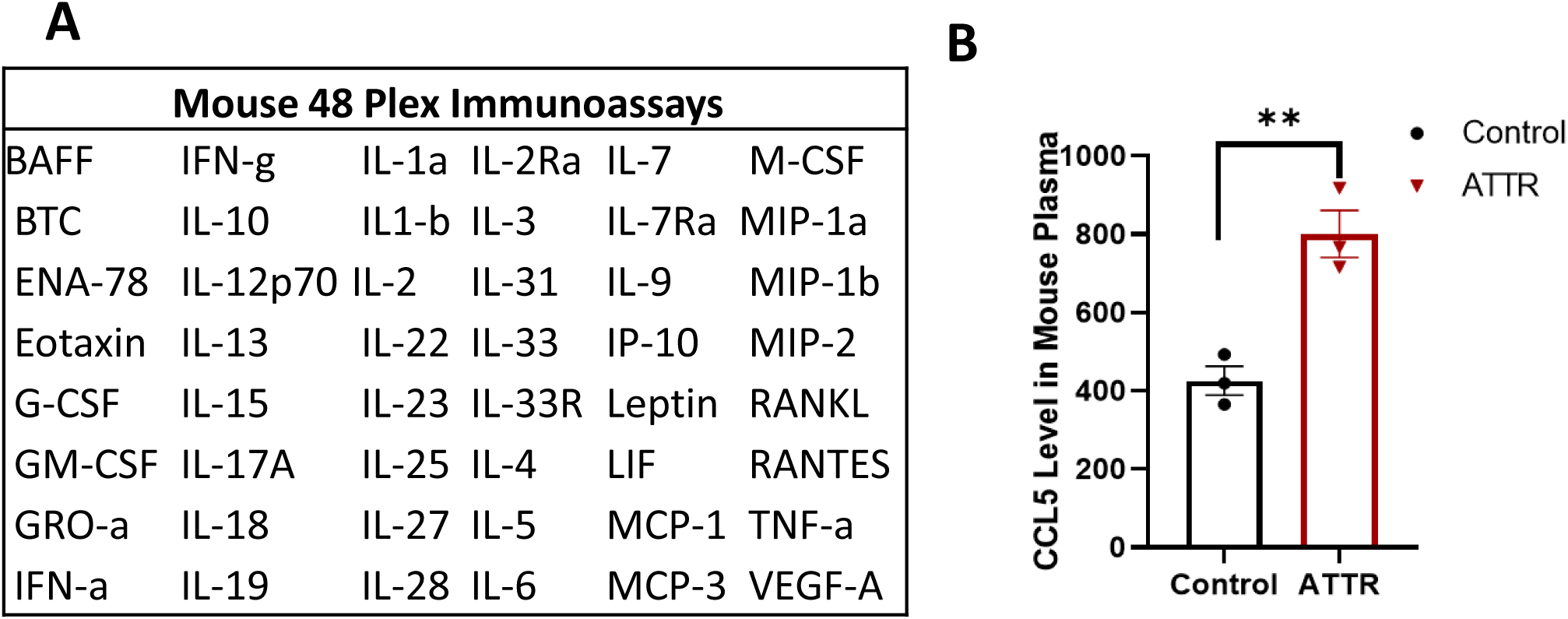
Elevated CCL5 in ATTR Murine Serum Post Injection 3 Months. **(A)** Mouse 48 Plex (Procarta) luminance-based immunoassay to check 48 common inflammatory cytokines in mouse plasma; **(B)** Averaged median fluorescence intensity (MFI) across duplicates and four control beads (Radix BioSolutions, Chex1-Chex4) were utilized for the different steps of the assay. Quantitative analysis of the MFI showed an increased level of CCL5 (MFI ATTR: 801 ± 105; Control: 426± 64; p=0.0061, N=3 mice/group) in ATTR murine serum compared with controls. n=3 mice/group; Data are presented as mean ± SEM. **P < 0.01 by nonparametric T-test.

## Discussion

Recent advancements in gene editing with CRISPR-Cas9 suggest that newer models may overcome the previous limitations, such as low human TTR expression, mild tissue-specific amyloid deposition, by allowing more precise replication of the human disease phenotype in mice [55]. Our study employed a TTR-KO mouse model over a wild-type to specifically address the biophysical and biochemical challenge posed by murine TTR, which naturally forms a stable human-mouse heterotetrameric, thereby impeding the aggregation and deposition of unstable human TTR [56, 57]. By leveraging the TTR-KO model, we aimed to enhance the expression of human TTR, leading to significant cardiac amyloid deposition and elevated circulating TTR levels in our humanized V122I ATTR mouse model. This approach was chosen to faithfully mimic the human condition, particularly in the context of age-related ATTR risk, by using older mice and employing advanced gene delivery and editing techniques to replicate human TTR production and achieve high transduction rates in the liver. The combination of CRISPR/Cas9 gene editing with AAV gene delivery not only overcomes previous limitations of rodent ATTR models, such as low serum TTR levels and lack of cardiac amyloid deposition but also enables our model to serve as a powerful tool for in-depth mechanistic studies of ATTR. The observed preservation of cardiac function despite amyloid deposition may reflect a need for a larger sample size and use younger mice with a longer follow-up to fully understand the impact of amyloidosis on cardiac function, prompting further studies with extended durations to assess the progression of cardiac alterations over time. Our ATTR mouse model could potentially serve as a preclinical pharmacogenomic model of novel therapeutics, which will undoubtedly improve outcomes for ATTR patients.

The precise molecular mechanisms underlying ATTR-CM remain unidentified. Our study uncovered a potential link between inflammation and ATTR-CM in which we identified upregulation of inflammatory signaling pathways through transcriptomic analysis of RNA sequencing for the ATTR murine hearts. Our data suggested that amyloid fibril deposition may alter the cardiac microenvironment, disturb the homeostasis, and trigger a cascade of the immune response, which we observed the upregulation of chemical carcinogenesis-reactive oxygen species, cellular senescence and platelet activation pathway changes in ATTR. This process can further exacerbate the activation of inflammatory cells, and the release of cytokines and chemokines leading to heart dysfunction, which we identified the T cell, B cell receptor signaling, leukocyte transendothelial migration, TNF signaling were upregulated, along with the increased expression of CXCL1, CXCL2, CXCL3 and CCL20 in ATTR murine hearts. Inflammatory cells can, in turn, contribute to impaired lysosomal function, further exacerbating the accumulation of precursor proteins in the heart tissue and inextricably linked to autophagy dysfunction, which we discovered the enhanced proteasome and apoptosis in ATTR hearts. Our findings suggest that the efficacy of certain anti-inflammatory medications, like diflunisal and corticosteroids, in treating cardiac amyloidosis could be due to their role in reducing inflammation and improving clinical outcomes [58, 59]. Our research indicates that chemokine-based treatment against inflammation holds potential as a therapeutic strategy for managing ATTR-CM.

Additionally, our findings demonstrated alterations in calcium levels and contractility in the V122I ATTR mouse model, potentially enhancing our comprehension of the origins of arrhythmias observed in ATTR-CM patients. This includes heightened cardiac muscle contraction, elevated calcium signaling, and calcium reabsorption influenced by endocrine and other factors. Intriguingly, our data suggested an upregulation of pathways associated with cardiac remodeling, indicating early changes in cardiac structure that might not be detectable via echocardiography. The MAPK pathway involved in cell growth, differentiation, and stress response, contributes to pathological cardiac remodeling and heart failure through its subfamilies, such as ERK (extracellular signal-regulated kinase), JNK (c-Jun N-terminal kinase), and p38 MAPK [60]. The JAK-STAT signaling pathway transduces signals from the cell surface to the nucleus, modulating gene expression in response to various stressors and is also implicated in cardiac hypertrophy, remodeling, and ischemia/reperfusion-induced dysfunction [61–63]. Our study provided the evidence linking MAPK and JAK-STAT pathways to ATTR pathogenesis, understanding these signaling mechanisms may offer insights into novel therapeutic targets for ATTR-related cardiac conditions.

The role of CCL and CXCL chemokine families in the pathogenesis of cardiovascular disease, particularly heart failure is complex and multifaceted. These chemokines play crucial roles in mediating inflammation, fibrosis, and immune cell recruitment, which are key processes in the development and progression of these conditions. Chemokines such as CCL2 and CXCL8 are significantly induced in infarcted hearts, contributing to leukocyte trafficking, inflammation, and subsequent cardiac remodeling. These actions underline the potential of targeting chemokines for therapeutic interventions in myocardial infarction and heart failure [64]. The chemokine receptor CXCR3 and its ligands (e.g., CXCL9, CXCL10) are implicated in various CVDs, including heart failure. These chemokines are involved in immune cell recruitment and vascular function, suggesting a complex role in CVD pathogenesis and potential as biomarkers for heart failure [65]. Chemokines like CCL2, CXCL12, and CXCL10 are involved in myocardial fibrosis [66]. The expression of chemokines such as CXCL9 and CXCL10 is associated with the intensity of myocarditis in Chagas cardiomyopathy, indicating their role in inflammatory cell migration to the myocardium [67].

Our data identified inflammation as a potential exacerbating factor in ATTR-CM, shed light that CCL and CXCL families may contribute to the pathogenesis of ATTR-CM. Specifically, we observed the significant elevations in CCL5, which also known as RANTES (Regulated on Activation, Normal T cell Expressed and Secreted) and plays an active role in recruiting a variety of leukocytes into inflammatory sites including T cells, macrophages, eosinophils, and basophils. Coordinating with certain cytokines that are released by T cells such as IL-2 and IFN-, CCL5 also induces the activation and proliferation of natural killer cells to generate C-C chemokine-activated killer cells [68]. This suggests CCL5 mediated inflammation may confer the ATTR-CM pathogenesis, pointing to the importance of further *in vivo* studies to decipher the detailed mechanism how inflammation modulates the ATTR-CM.

This study has several limitations. First, we aim to increase the sample size and improve our ability to monitor early cardiac function changes by incorporating a 3D echocardiogram to calculate relative wall thickness (RWT) compared to LV mass using IVSd, LVPWd, and LVIDd measurements. If RWT significantly differs from controls, this could indicate early cardiac remodeling. Additionally, we plan to validate our findings using ATTR patient biopsies. Our team includes researchers from two of the leading clinical and research centers in amyloidosis in the U.S.—the Stanford Amyloid Center and the Brigham and Women’s Hospital Cardiac Amyloid Program. We are establishing an ATTR biobank, which will include human cardiac tissue and blood samples from patients with ATTR-CM, non-amyloid heart failure, and healthy controls, for future validation studies. To further enhance and optimize our mouse model, we intend to isolate patient-specific amyloid fibrils from ATTR-CM heart biopsies and assess whether these human-derived fibrils exacerbate cardiac amyloid deposition in our V122I ATTR mouse model.

In summary, our humanized V122I ATTR mouse model provides a promising and invaluable research tool for preclinical therapeutics development for ATTR patients. Our study suggests that inflammation may play an important role in ATTR-CM pathogenesis. Further *in vivo* studies are needed to dissect detailed mechanisms of how inflammation modulates ATTR-CM.

## Conclusion

In this study, we generated a novel humanized V122I ATTR mouse model to overcome the previous drawbacks of the transgenic mouse models onto a murine TTR-null background [69, 70]. The murine TTR gene had been silenced and stabilizes the circulating mutant protein to in vitro urea denaturation. Our V122I ATTR mouse showed significantly elevated circulating human TTR levels with Congophilic amyloid deposits in the murine heart and kidneys. Furthermore, both immature and mature amyloid fibrils were observed in ATTR murine hearts by TEM. The comprehensive transcriptomic analysis of the ATTR murine hearts revealed the upregulation of inflammation in ATTR, specifically CXCL- and CCL-chemokines were identified and further confirmed a significantly increased level of CCL5 at post translations level in mouse plasma. Taken together, we developed a humanized mouse model to replicate the multisystem complexity and clinical diversity associated with V122I ATTR which will allow for further *in vivo* mechanistic studies. Our data suggested that CXCL- and CCL-chemokine may contribute to the pathogenesis of ATTR-CM. Our study provided pivotal evidence to uncover the molecular signature of inflammation in ATTR-CM and the chemokine-based therapeutic against inflammation may be beneficial to ATTR-CM. Further *in vivo* studies are needed to decipher the precise interactions between inflammation and ATTR.

## Acknowledgments

We thank Drs. Ronglih Liao,Joseph Woo, Hiroki Kitakata and Gracia Fahed for critical suggestions on the manuscript; Drs. Dean Felsher and Joanna E. Lilienthal for generous TRAM funding support and mentorship; Nixuan Cai, John Isaiah Jimenez and Alessandro Evangelisti for skillful assistance with experiments related to animal breeding, genotyping, western blot, ELISA, histology studies; Alicia Wu and Leavy Hu assistant with Congo Red image acquisition and analysis; Dr. Alokkumar Jha for RNA-seq transcriptomic analysis; Dr. Nerea Zabaleta and her members of Harvard Medical School MEEI/ SERI Gene Transfer Vector Core for AAV vector production; John J. Perrino from Stanford Transmission Electron Microscope Facilities for TEM technical support; Iris Herschmann and Dr. Holden Terry Maecker from Stanford Human Immune Monitoring Center (HIMC) for immunoassay and technical support. Work in the laboratory of Dr. Kevin Alexander is supported by the funding from Stanford Department of Medicine, the Harold Amos Medical Faculty Development Award from the American Heart Association, Stanford CTSA KL2 Mentored Career Development Award from the National Institutes of Health. This work was also supported by the Stanford Translational Research and Applied Medicine (TRAM) postdoc pilot grant (2021-2023).

## Contributions

X.W. and K.M.A. conceived the project. K.M.A. supervised the project. X.W., N.C., I.J., and A.E. designed the experiments. X.W., N.C., I.J., and A.E. performed the experiments. X.W., N.C., I.J., A.E., and A.J., analyzed the data. X.W., N.C., I.J., and K.M.A. wrote the manuscript. X.W., N.C., I.J., A.E., A.J., H.K., J.W., R.L., and K.M.A reviewed the manuscript. All authors read, corrected, and approved the manuscript.

## Ethics declarations

NA

## Nonstandard Abbreviations and Acronyms

ATTR: Transthyretin cardiac amyloidosis
ATTR-CM: Transthyretin Cardiomyopathy
TTR-KO: TTR knockout
TEM: Transmission Electron Microscopy
AL: Immunoglobulin light chain amyloid
TTR: Transthyretin
V122I: TTR mutation that substitutes valine for isoleucine at position 122
wtATTR-CM: Wild-type transthyretin cardiomyopathy
HFpEF: Heart failure with preserved ejection fraction
TAVR: Transcatheter aortic valve replacement
PIK3C3: A protein with a central role in autophagy
PVDF: Polyvinylidene difluoride
ELISA: Human Prealbumin Enzyme-linked Immunosorbent Assay
BSA: Bovine serum albumin
LV: Left ventricular
LVAWd: LV anterior wall end-diastolic thickness
LVAWs: LV anterior wall end-systolic thickness
LVIDd: LV internal dimension at end-diastole
LVIDs: LV internal dimension at end-systole
LVPWd: LV posterior wall end-diastolic thickness
LVPWs: LV posterior wall end-systolic thickness
IVSd: Interventricular septum thickness end diastole
HR: Heart rates
EF: Ejection fraction
FS: Fractional shortening
OCT: Optimal cutting temperature
EtOH: Ethanol
GAPDH: Glyceraldehyde 3-phosphate dehydrogenase
GO: Gene Ontology
GSEA: Gene Set Enrichment Analysis
HIMC: Stanford Human Immune Monitoring Center
MFI: Median fluorescence intensity
AAV: Adeno-associated Virus
GTVC: Gene Transfer Vector Core
HW/BW: Heart-to-body weight
LW/BW: Lung-to-body weight ratio
HW/TL: Heart weight to tibial length
LW/TL: Lung weight to tibial length
ECM: Cardiac extracellular matrix
DEGs: Differentially expressed genes
RANTES: Regulated on Activation, Normal T cell Expressed and Secreted

