## Supplemental Figures 1-6 for "The role of inflammation in a humanized mouse model of transthyretin cardiac amyloidosis"

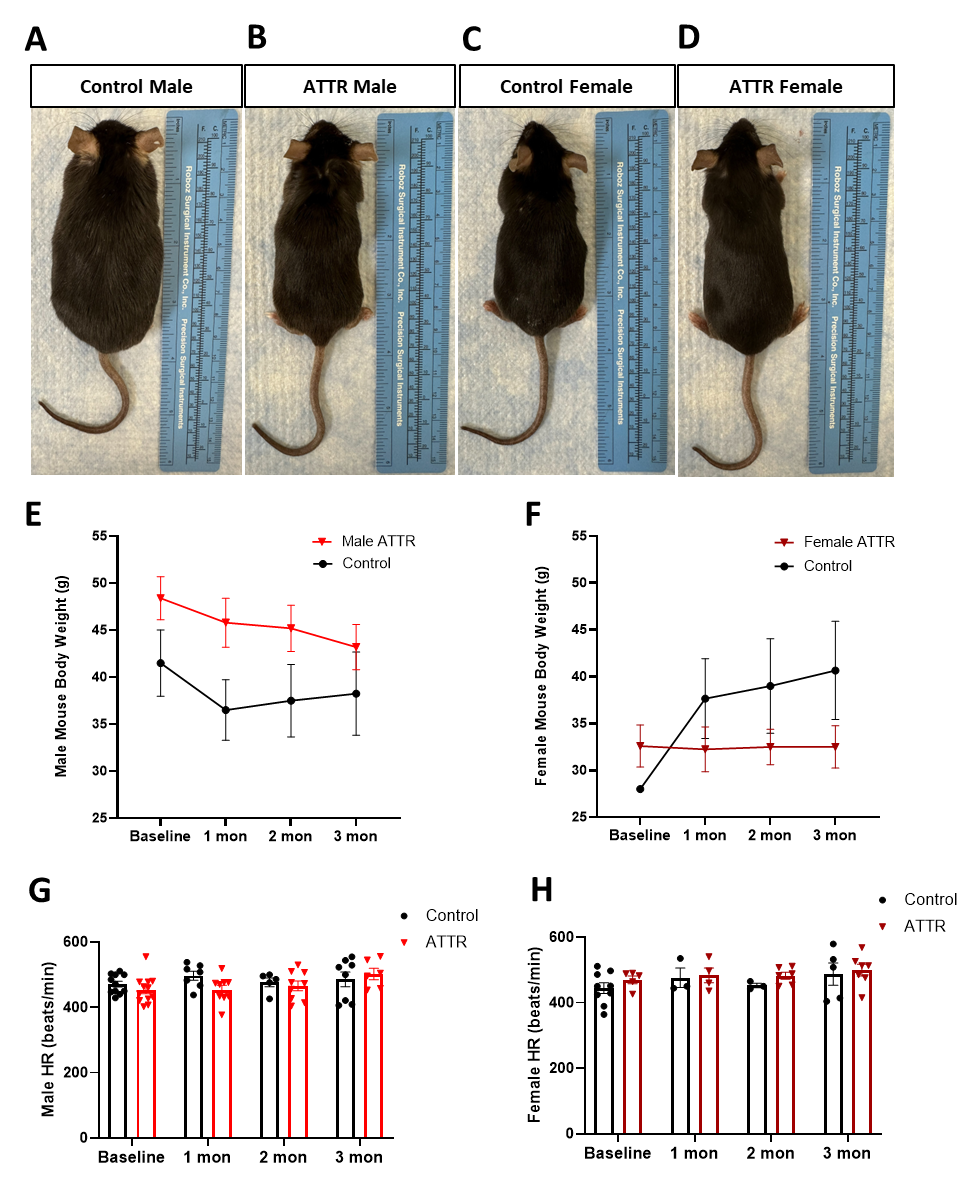

**Supplemental Figures**

**Supplemental Figure 1: Morphology and Body Weight Changes of the ATTR Mouse**

**(A-D)** Representative images of male, female ATTR, and control mice post-injection 3 months. **(E-F)** Quantitative analysis of male and female body weight change during time course. There were significant body weight changes between male and female ATTR mice compared with controls during post-injection time. N=3-5 mice/group; **(G-H)** Quantitative heart rate (HR) analysis in male and female mice post injection for 3 months; Data are presented as mean ± SEM—statistically analysis by two-way ANOVA.

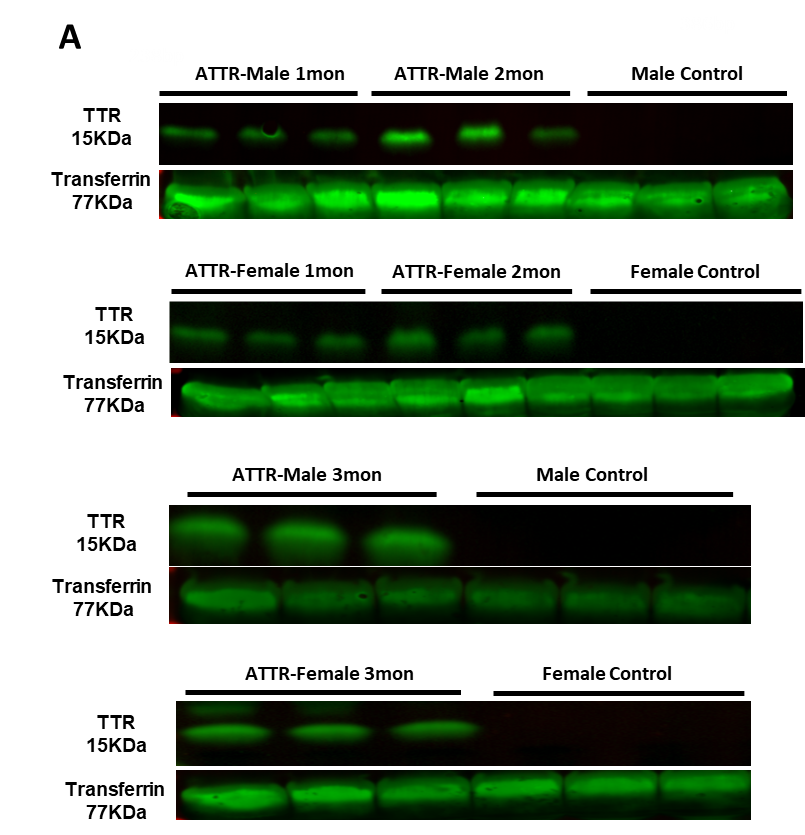

**Supplemental Figure 2: Human TTR Expression Level in ATTR Mouse by Western Blots.**

(A) Raw images of western blots of human TTR level in serum from male and female ATTR and control mice during time course 1-,2-,3-month, transferrin was utilized as loading control. (1st antibody Anti-human TTR Flag 1:1000; 2nd antibody Goat anti-Rabbit 1:10,000).

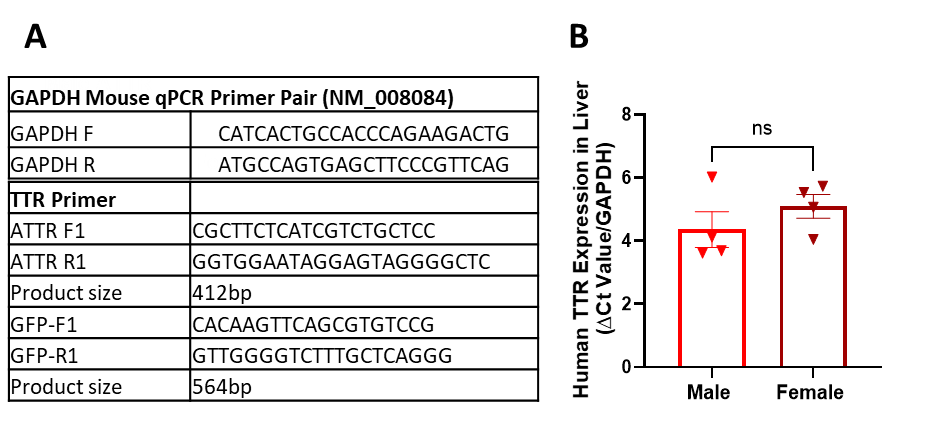

**Supplemental Figure 3: Human TTR Expression Efficiency in Liver Post Injection**

(A) RT-qPCR for human TTR expression level in male and female ATTR mice 3 months post-injection. RNA was extracted from murine livers of ATTR and control mice. (B)The human TTR primers sequence is used in electrophoresis and RT-qPCR. GAPDH primer sequences were used as an internal control.

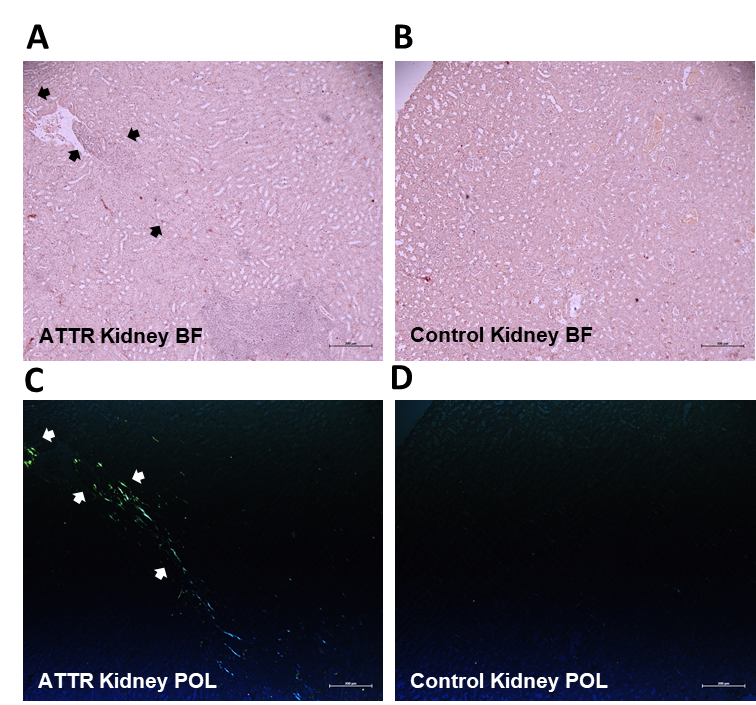

**Supplemental Figure 4. Renal Amyloid Deposition in ATTR Mouse by Congo Red Staining.**

(A-B) Representative images of Congo Red staining for male ATTR kidney biopsy and control mouse under bright field (BF) Original magnification, 5x. The black arrows indicated the amyloid fibrils were stained as red. (C-D) Representative images of Congo Red staining male ATTR kidney biopsy and control mouse under polarized light (POL) imaging. The white arrows indicated the amyloid fibrils were shown as apple green under polarized view. Original magnification, 5x.

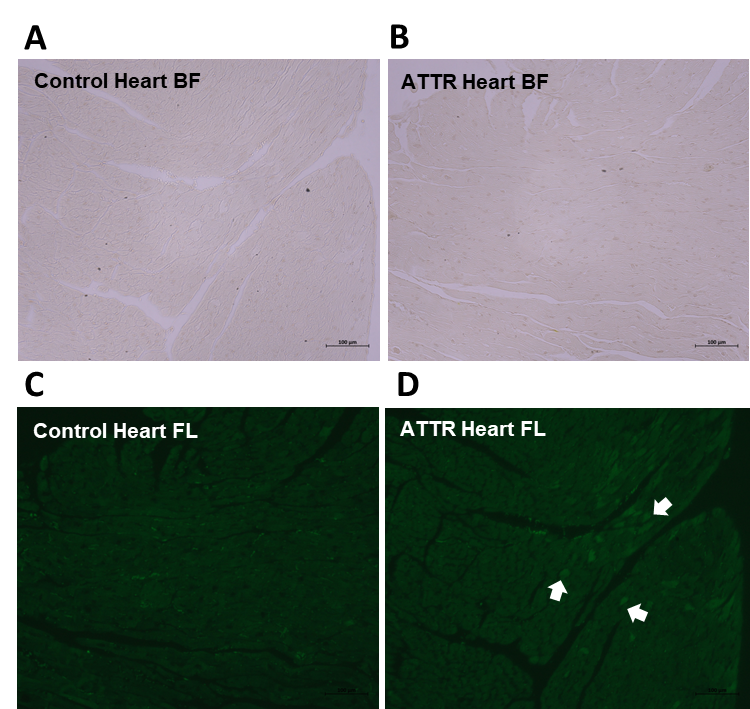

**Supplemental Figure 5: Cardiac Amyloid Deposition in ATTR Mouse By Thioflavin T Staining.**

(A-B) Representative images of Thioflavin T staining for male ATTR mouse and control mouse hearts under bright field (BF). Original magnification, 10x. (C-D) Representative pictures of Thioflavin T staining (ThT) for male ATTR mouse and control mouse hearts under fluorescence imaging (FL, Excitation 450nm/ Emission 488nm). Original magnification, 10x. The white arrows indicated the amyloid fibrils were shown as green fluorescence.

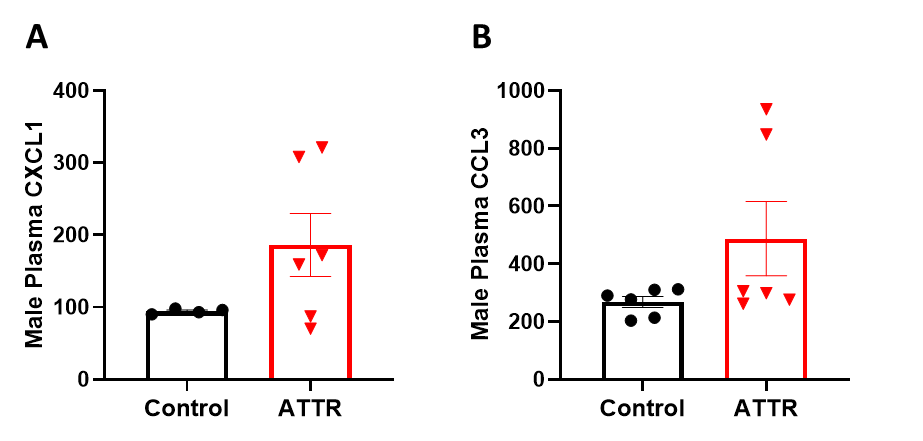

**Supplemental Figure 6. Mouse 48 Plex (Procarta) luminance-based immunoassay to check inflammatory cytokines in mouse plasma.**

(A-D) Quantitative analysis of CXCL1 and CCL3 in male plasma from ATTR and control mice. Averaged median fluorescence intensity (MFI) across duplicates was analyzed. N=3 mice/group; Data are presented as mean ± SEM—statistical analysis by nonparametric T-test.
